# Membrane-catalyzed aggregation of islet amyloid polypeptide is dominated by secondary nucleation

**DOI:** 10.1101/2022.02.04.479144

**Authors:** Barend O.W. Elenbaas, Lucie Khemtemourian, J. Antoinette Killian, Tessa Sinnige

## Abstract

Type II diabetes is characterized by the loss of pancreatic β-cells. This loss is thought to be a consequence of membrane disruption, caused by the aggregation of islet amyloid polypeptide (IAPP) into amyloid fibrils. However, the molecular mechanisms of IAPP aggregation in the presence of membranes have remained unclear. Here, we use kinetic analysis to elucidate the aggregation mechanism of IAPP in the presence of mixed zwitterionic and anionic lipid membranes. The results converge to a model in which aggregation on the membrane is strongly dominated by secondary nucleation, i.e. the formation of new nuclei on the surface of existing fibrils. The critical nucleus consists of a single IAPP molecule, and anionic lipids catalyze both primary and secondary nucleation, but not elongation. The fact that anionic lipids promote secondary nucleation implies that these events take place at the interface between the membrane and existing fibrils, demonstrating that fibril growth occurs at least to some extent on the membrane surface. These new insights into the mechanism of IAPP aggregation on membranes may help to understand IAPP toxicity and will be important for the development of therapeutics to prevent β-cell death in type II diabetes.

## Introduction

Type II diabetes is a major health problem affecting about 500 million people worldwide^1,2^. This disease is characterized by insulin resistance and dysfunction of the insulin-producing β-cells in the pancreas^3,4^. A key histopathological hallmark of type II diabetes is the presence of fibrillar deposits formed by islet amyloid polypeptide (IAPP), a 37 amino acid-long hormone with various functions that is co-secreted with insulin from the β-cells. The process of fibril formation by IAPP is thought to damage the membranes of the β-cells, ultimately causing cell death^5,6^.

The mechanism by which IAPP aggregation causes toxicity is not entirely understood. One hypothesis is that IAPP forms discrete pores in the membrane^7,8^. Soluble oligomers have also been found to cause damage to synthetic lipid bilayers and cell membranes^9–11^. Fibril growth has been suggested as another mechanism of causing membrane damage, involving deformation of vesicles^12^ as well as the extraction of lipids from the bilayer^13^.

The aggregation of IAPP follows the typical nucleation-dependent polymerization process that is characteristic for amyloid formation. The strongly sigmoidal aggregation curves obtained for IAPP under solution conditions have been attributed to secondary nucleation, an auto-catalytic process by which nucleation events are catalyzed on the surface of existing fibrils^14–16^. In studies using synthetic human IAPP, the rates of fibril formation were found to be relatively independent of the IAPP monomer concentration, suggesting that the peptide exists as pre-formed condensates (also referred to as oligomers or micelles)^14,17^. A different type of IAPP preparation from a recombinant source recently allowed for a quantitative kinetic analysis of Thioflavin T (ThT) data, which confirmed the important contribution of secondary nucleation^16^.

Monomeric IAPP interacts with membranes by adopting a partial α-helical structure^18–20^. The α-helical signature disappears upon aggregation of IAPP, when a β-sheet structure becomes apparent^19,20^. It has been suggested that the α-helical structure is off-pathway to fibril formation, given the faster aggregation of IAPP variants unable to form a helix^21^. In spite of this, membranes containing anionic lipids have been shown to accelerate IAPP fibril formation^20,22–24^. The anionic head groups likely interact with the positively charged residues in the N-terminal half of the peptide, yet how this interaction promotes IAPP aggregation remains unclear.

A detailed characterization of IAPP fibrillization in the presence of lipids is essential to make progress in understanding how this process relates to membrane damage. However, mechanistic insights into IAPP aggregation have so far been restricted to solution conditions. Here, we use kinetic analysis to establish a molecular mechanism for IAPP fibril formation catalyzed by membranes containing anionic lipids. Global fitting of kinetic data to mathematical models is a powerful tool to establish the microscopic steps of fibril formation and to quantify the associated rates^25–27^. Despite the complexity of this system, we show that a relatively simple aggregation model is sufficient to explain the membrane-catalyzed fibril formation of IAPP.

## Materials and Methods

### Materials

1,2-dioleoyl-*sn*-glycero-3-phosphocholine (DOPC) and 1,2-dioleoyl-*sn*-glycero-3-phospho-L-serine (DOPS) were acquired from Avanti Polar Lipids. Islet amyloid polypeptide (IAPP) was purchased from Bachem (Amylin (human) trifluoroacetate salt, Lot no. 1000026803). All other chemicals were purchased from Sigma-Aldrich.

### IAPP preparation

IAPP from the manufacturer was dissolved in hexafluoro-2-propanol to a concentration of 1 mM and the solution was left on the bench at room temperature for 2 h. Aliquots were made by dividing the stock in separate microcentrifuge tubes which were placed in high vacuum for 1 h. The aliquots were stored at -80 °C and dissolved in milliQ water at 160 µM prior to use.

### Seed preparation

Monomeric IAPP was dissolved in 10 mM Tris-HCl, 100 mM NaCl pH 7.4 at 50 µM final concentration and allowed to aggregate at room temperature for approximately 24 h. Subsequently, the formed fibrils were sonicated in a bath sonicator (Branson, 3800) for 2 min and the resulting seeds were used within 15 min of sonication.

### Vesicle preparation

Unless specified otherwise, a membrane composition of 7:3 DOPC/DOPS (mol:mol) was used. Stock solutions of DOPC and DOPS were made in chloroform and the lipid concentrations were determined using a Rouser assay^28^. The appropriate volumes of each were mixed together in chloroform and dried under a nitrogen gas flow in a 42 °C water bath to evaporate the solvent. The samples were placed in a desiccator for at least half an hour to remove any residual solvent. The lipid film was then resuspended in 10 mM Tris-HCl pH 7.4, 100 mM NaCl to allow for the spontaneous formation of multilamellar vesicles (MLVs). These were left on the bench at room temperature for 1 h with gentle shaking every 10 min to completely suspend the lipid film. The MLVs were subjugated to 10 freeze thaw cycles by alternatingly placing them in ethanol cooled by dry ice pellets, and lukewarm water. To create large unilamellar vesicles (LUVs), the MLVs were extruded through a 200 nm pore filter (Anotop 10, Whatman, Maidstone, U.K) using syringes. This was done 10 times back and forth, followed by one more passage to ensure that the vesicles ended on the opposite side from where they started. The final lipid concentration of the LUV suspension was determined again using a Rouser assay^28^. LUVs were stored at 4 °C and used within one week after preparation.

### Thioflavin T assay

All experiments were performed in triplicate, in 200 µL volumes with final buffer conditions of 10 mM Tris-HCl pH 7.4, 100 mM NaCl and 10 µM ThT. The plates were kept on ice during the entire pipetting procedure up to the moment they had to be placed in the plate reader. The IAPP solution was also kept on ice and added last. The ThT-assays were performed in a climate room at 20 °C on a CLARIOstar plus (BMG Labtech) plate reader. A transparent cover sticker (Viewseal sealer, Greiner) was used to prevent evaporation during the measurement. Before the first measurement, the 96 well, flat bottom plates (CELLSTAR black, Greiner) were shaken once for 20 seconds at 500 rpm, linearly. The used excitation/emission settings were 430-15 nm/535-15 nm with 482.5 nm dichroic mirror setting and with a 3 mm orbital averaging of 53 flashes per well. The data shown for the different IAPP:lipid ratios and DOPC:DOPS ratios are representative of at least two independent datasets.

### Global fitting

The AmyloFit webserver was used for global fitting of the kinetic data^27^. The raw ThT fluorescence data were transformed into the required format as described in the protocols provided with AmyloFit. The equation used to fit the data to a secondary nucleation dominated mechanism is^26,27^:

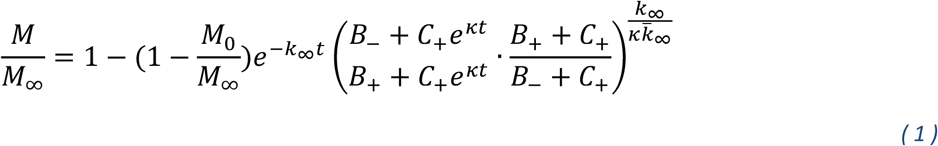

using:

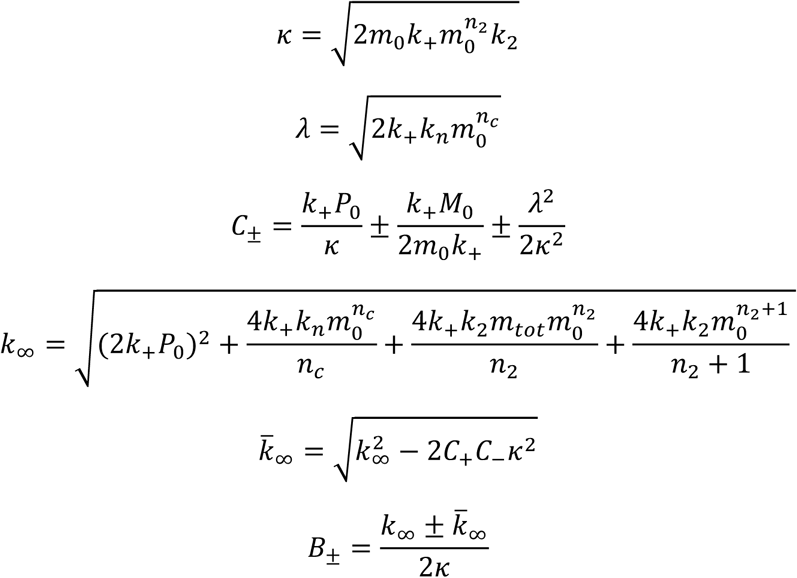

where *m*_*0*_ is the initial monomer concentration, *M* the fibril mass concentration, *P* the number concentration of fibrils, *n*_*c*_ and *n*_*2*_ the reaction orders of primary and secondary nucleation, and *k*_*n*_, *k*_*2*_ and *k*_*+*_ the rate constants for primary nucleation, secondary nucleation and elongation, respectively.

When fitting the data, *n*_*c*_ and *n*_*2*_ were fixed to either 1 or 2 as specified in the text, and the combined rate constants *k*_*+*_*k*_*n*_ and *k*_*+*_*k*_*2*_ were fitted as global parameters.

### Seeded IAPP aggregation

The relationship between the initial gradient and the elongation rate constant for highly seeded reactions is described by the following equation^27^:

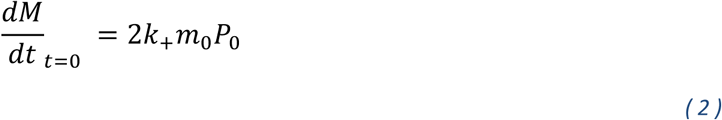

where *dM/dt*_*t=0*_ is the initial gradient of the fibril mass concentration *M, k*_*+*_ is the elongation rate constant, *m*_*0*_ is the initial monomer concentration and *P*_*0*_ is the initial number of seed fibrils. *m*_*0*_ and *P*_*0*_ are the same for the reactions with 30% and 70% DOPS. Thus, the initial gradient for these seeded reactions can be used to compare the *k*_*+*_ for both reactions.

The half-times of the seeded reactions were plotted against the logarithm of the seed concentrations, which should scale linearly in case of a mechanism dominated by secondary nucleation^29^.

### Linear regression

Linear regressions to determine the scaling exponent were performed in Graphpad Prism version 8.0.2, using the default analysis settings. The scaling exponent *γ* corresponds to the slope of the double-log plot according to the equation^27^:

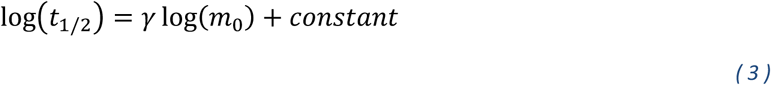

where the base of the log is the same on both sides but can have any value.

## Results

### IAPP aggregation kinetics in the presence of membranes

In order to establish the contribution of lipid membranes to the aggregation kinetics of IAPP, we first performed the aggregation assay in the absence of membranes. Under these conditions, the kinetics of IAPP aggregation are not concentration-dependent as can be seen from the overlap between the normalized ThT curves (**Fig. 1A, Fig. S1A**). These results are in agreement with previous studies using synthetic peptide that attributed this behavior to the existence of IAPP condensates or micelles, resulting in a constant aggregation rate that is independent of the bulk monomer concentration^14,17^. Similar findings were recently reported for another amyloid-forming protein, α-synuclein, in its phase-separated state^30^.

**Figure 1.**
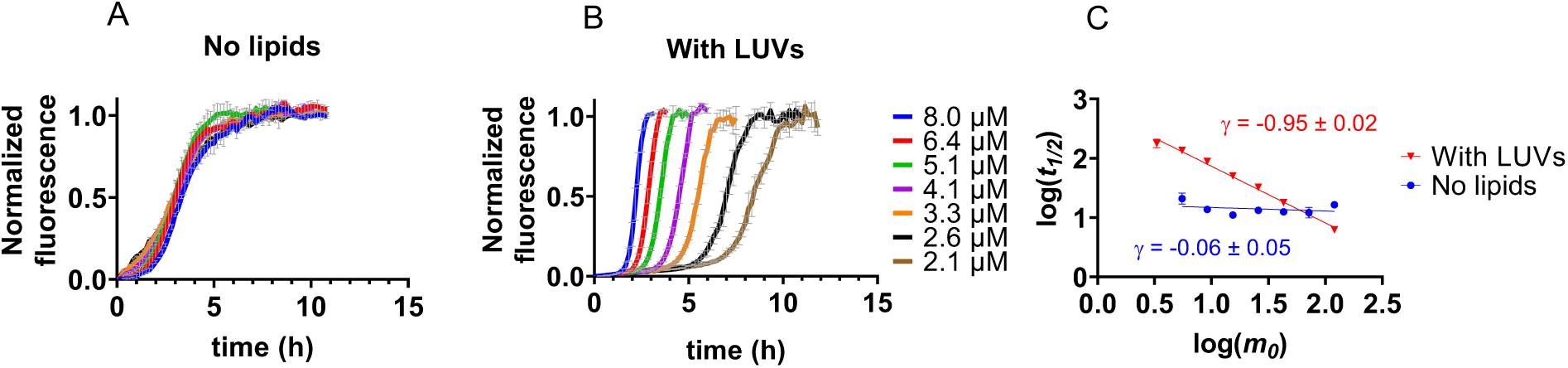
Concentration dependence of IAPP aggregation in the absence and presence of lipid membranes. (A, B) Normalized ThT fluorescence for IAPP aggregation in the absence of membranes (A) and in the presence of DOPC/DOPS LUVs at a 1:100 IAPP:lipid molar ratio (B). The traces in (A) and (B) represent the average with standard deviation of three technical replicates. (C) Double log plot of the half-time (*t*_*1/2*_) versus the IAPP monomer concentration at t = 0 (*m*_*0*_). Scaling exponents (*γ*, slope of the line through the datapoints) were derived by linear regression.

A different view emerges when looking at IAPP aggregation kinetics in the presence of large unilamellar vesicles (LUVs) composed of zwitterionic and anionic lipids (DOPC and DOPS, respectively) in a molar ratio of 7:3. At a peptide: total lipid molar ratio of 1:100, aggregation becomes faster with increasing IAPP monomer concentration (**Fig. 1B, Fig. S1B**). To assess the concentration dependence, a double-log plot was created, in which the logarithm of the initial monomer concentration (*m*_*0*_) is plotted against that of the half-time (timepoint at which half the maximum fluorescence intensity is reached, *t*_*1/2*_)^27^. For IAPP in the presence of LUVs, the datapoints lie on a straight line with a slope of approximately -1.0, which is the value of the scaling exponent (*γ*) (**Fig. 1C**). By contrast, in the absence of lipids, the scaling exponent of IAPP is close to 0, reflecting the lack of concentration dependence (**Fig. 1C**). Interestingly, the two lines intersect, showing that the addition of LUVs only accelerates the aggregation of IAPP above a monomer concentration of ∼ 6 µM.

### Global fitting of IAPP aggregation kinetics

The scaling exponent can be used to narrow down the possible aggregation mechanisms, which each have a unique dependence on the monomer concentration^27^. We used the online platform AmyloFit to fit the aggregation data of IAPP in the presence of LUVs to mathematical models representing these different mechanisms^27^. A scaling exponent of -1 may correspond to a simple mechanism consisting of nucleation and elongation, with a reaction order for nucleation (*n*_*c*_) of 2. However, this mechanism does not provide a satisfactory fit to our data (**Fig. 2A**). Another possibility is a mechanism incorporating secondary nucleation with a reaction order (*n*_*2*_) of 1, which does represent the data well (**Fig. 2B**).

**Figure 2.**
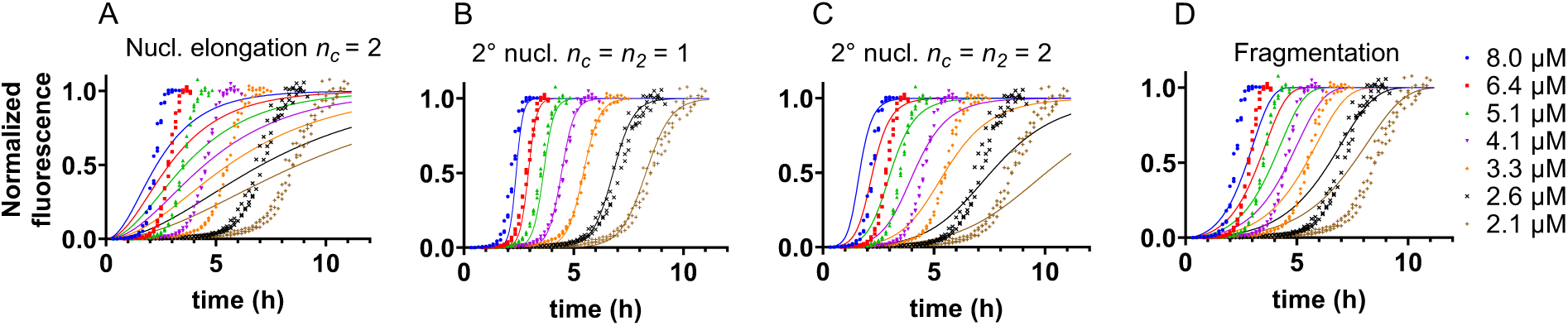
Global fitting of IAPP aggregation kinetics in the presence of LUVs. (A) Fits of ThT aggregation data to a nucleation elongation model with a reaction order for nucleation (*n*_*c*_) of 2. (B) Fits to a model in which secondary nucleation dominates and both reaction orders for primary (*n*_*c*_) and secondary nucleation (*n*_*2*_) are equal to 1. (C) Fits to a model in which secondary nucleation dominates and both reaction orders are equal to 2. (D) Fits to a fragmentation dominated model. The aggregation assay was performed in the presence of 7:3 DOPC:DOPS vesicles at a total IAPP:lipid molar ratio of 1:100.

Mechanisms with different scaling exponents, such as secondary nucleation with a reaction order of 2 (theoretically *γ* = -1.5) or fragmentation (*γ* = -0.5)^27^, do not fit our data as expected (**Fig. 2C, D**). More complex mechanisms like multi-step secondary nucleation or a combination of secondary nucleation and fragmentation do lead to reasonable fits (**Fig. S2A, B**). However, these mechanisms should display a curvature in the scaling exponent^27^, which we do not observe within the concentration range used in our experiments (**Fig. 1C**).

The best fit to the data (**Fig. 2B**) thus suggests that IAPP aggregation on lipid membranes is dominated by secondary nucleation with a reaction order of 1, corresponding to a critical nucleus size of a single monomer. The rate constants extracted from these fits reveal that secondary nucleation dominates over primary nucleation by many orders of magnitude (**Table 1**). Given the relatively minor contribution of primary nucleation, it is difficult to establish the reaction order of this process, with both values of 1 and 2 yielding similarly good fits to the datasets here presented (compare **Fig. 2B** with **Fig. S2C**). Primary and secondary nucleation are generally considered to be the same molecular conversion, and we favor the solution with both reaction orders being 1 as further detailed below.

**Table 1.**
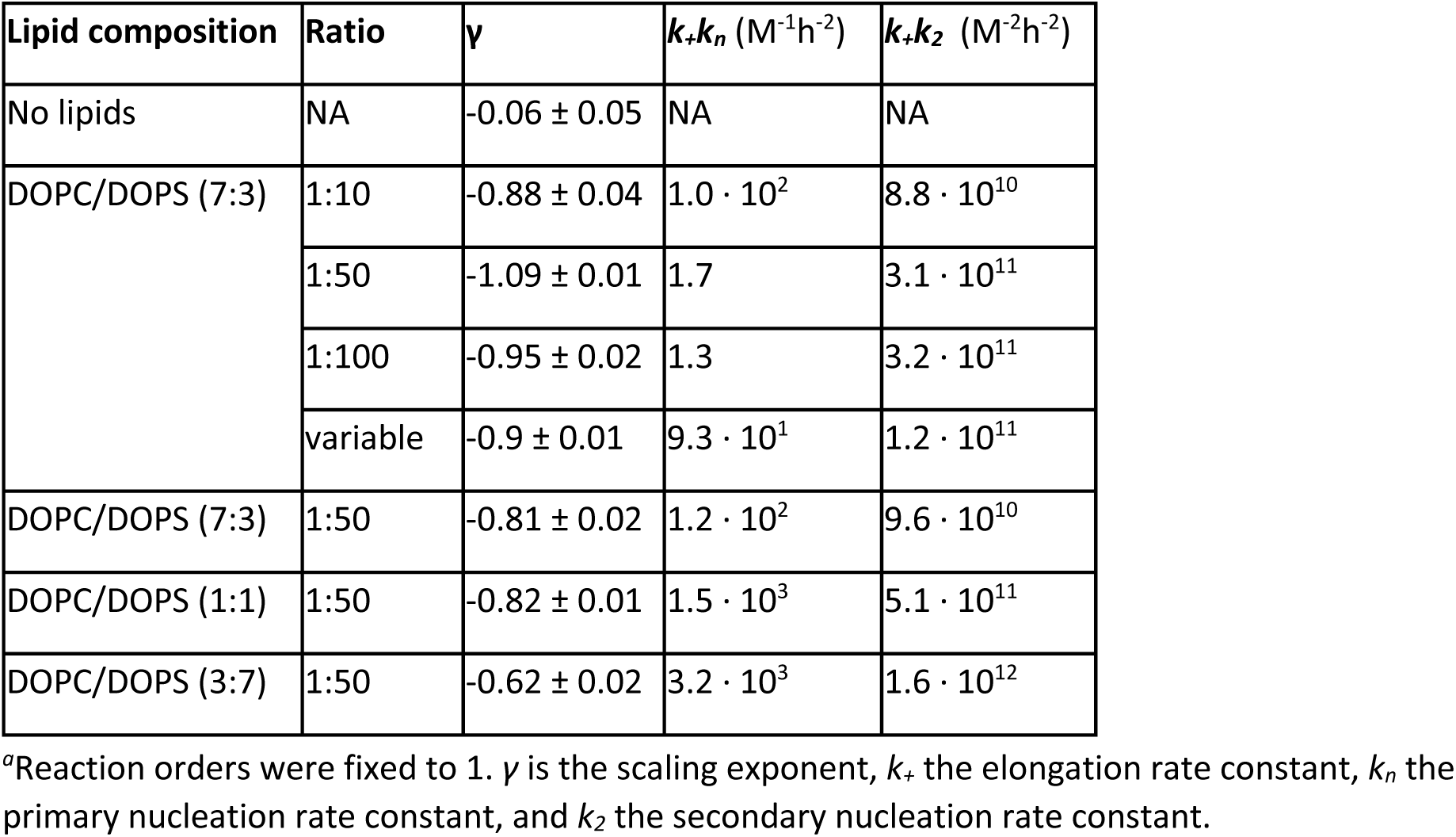
Values of the scaling exponents and rate constants derived from global fitting^*a*^

### IAPP aggregation kinetics are independent of the peptide:lipid ratio

If the critical nucleus size is indeed a monomer, the nucleation process should be independent of the local IAPP density on the membrane. To test this hypothesis, we performed the aggregation assay at lower IAPP:lipid ratios of 1:10 and 1:50, at which the IAPP density on the vesicles is expected to be higher (**Fig. 3, Fig. S3**). The scaling exponents are very similar (**Fig. 3A**), and the datasets fit well to the same model of secondary nucleation with a reaction order of 1 (**Fig. 3B, C, Fig. S4, Fig. S5**). Furthermore, the rate constants extracted from the fits to the 1:50 and 1:100 ratios are in close agreement (**Table 1**). Thus, the aggregation kinetics are essentially independent of the local peptide concentration on the vesicles. At a ratio of 1:10, we suspect that some IAPP that is not bound to the membrane aggregates in solution, and the scaling exponents and rate constants contain a minor contribution from this process.

**Figure 3.**
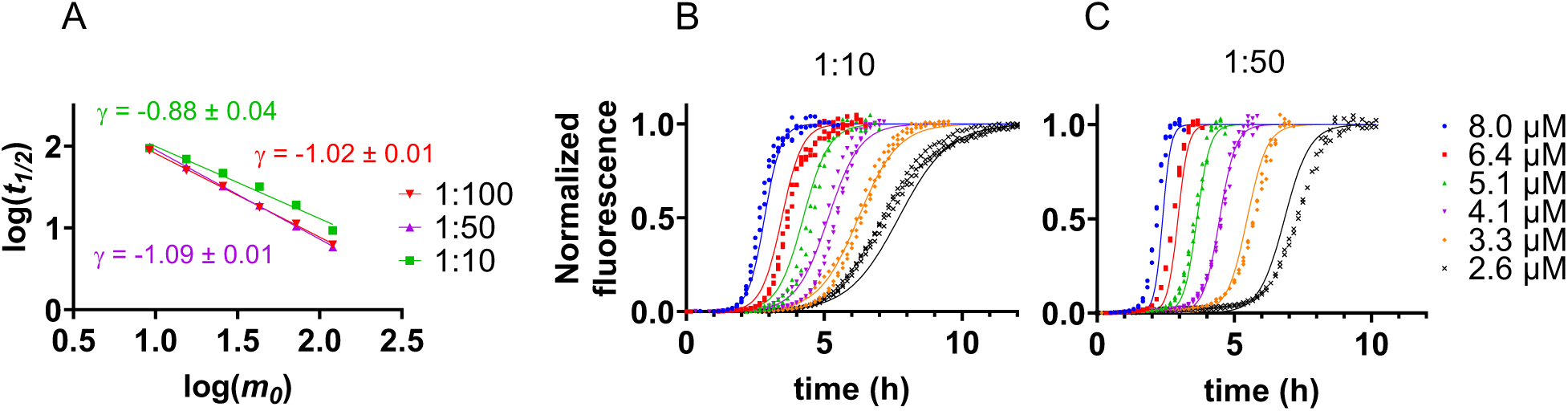
IAPP aggregation kinetics at varying peptide:lipid ratios. (A) Double-log plot with scaling exponents of aggregation assays performed at 1:100 (red), 1:50 (magenta), and 1:10 (green) IAPP:lipid ratios. A lipid composition of DOPC:DOPS 7:3 was used. (B) Fits of the 1:10 dataset to a model dominated by secondary nucleation with reaction orders for both nucleation processes equal to 1. (C) Fits of the 1:50 dataset to a model dominated by secondary nucleation with reaction orders for both nucleation processes equal to 1.

In a complementary experiment we used a constant concentration of lipids, which leads to increased crowding of IAPP on the vesicles with increasing IAPP concentration (**Fig. 4A, Fig. S6**). The scaling exponent and global fitting are again consistent with the same mechanism dominated by secondary nucleation with a reaction order of 1 (**Fig. 4B-D**). Under these experimental conditions, a reaction order of 1 is also the preferred solution for primary nucleation (compare blue and black lines in **Fig. 4C** and **Fig. 4D**). Altogether, our data indicate that a single IAPP monomer is able to undergo nucleation on the membrane.

**Figure 4.**
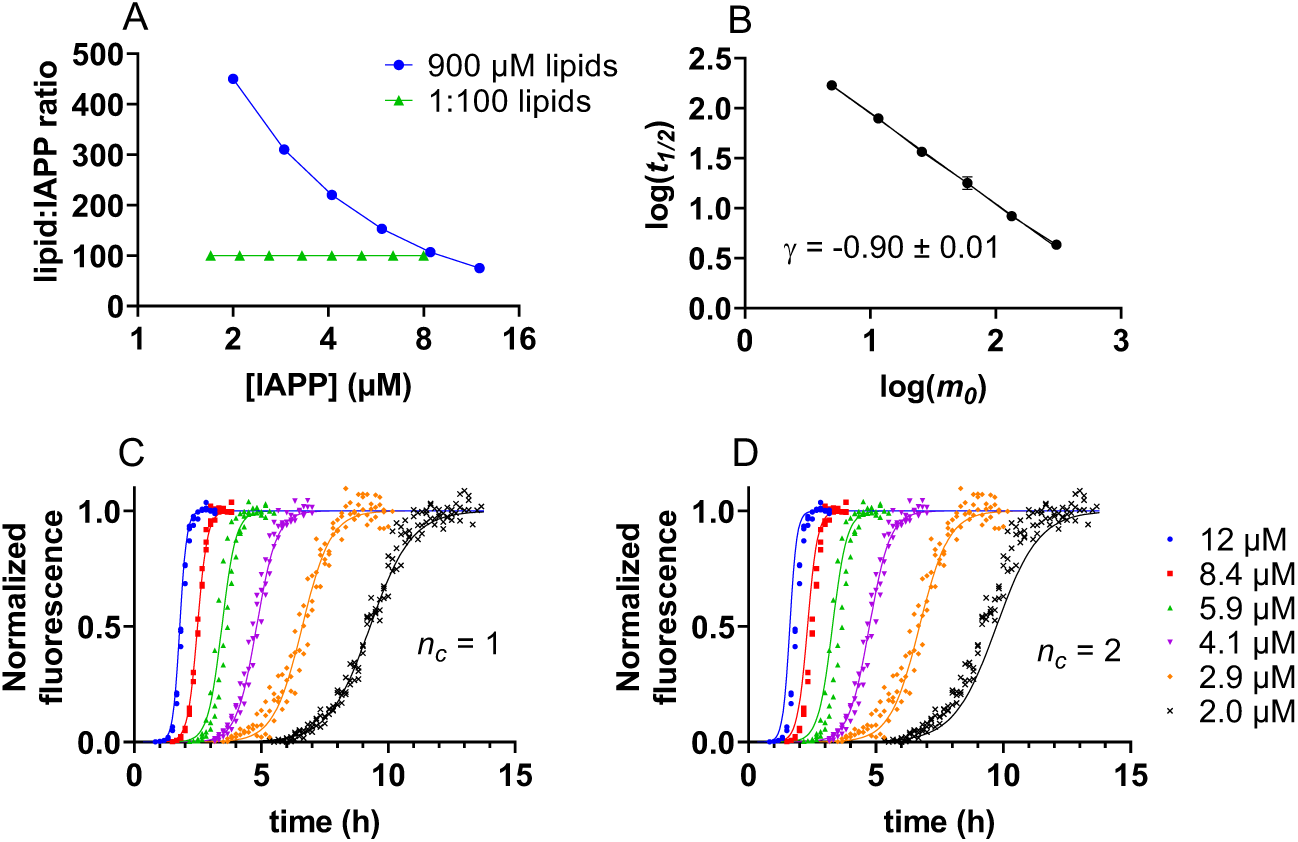
IAPP aggregation at a constant lipid concentration. (A) Graphical depiction of the number of lipids for each IAPP molecule at different experimental conditions: constant lipid concentration (blue) and constant ratio of 1:100 (green). (B) Scaling exponent derived from an IAPP aggregation assay performed at a constant lipid concentration of 900 µM for all IAPP concentrations. (C) ThT fluorescence for IAPP aggregation at a constant lipid concentration of 900 µM, fitted to a secondary nucleation dominated mechanism. Reaction orders for primary (*n*_*c*_) and secondary nucleation (*n*_*2*_) were both set to 1. (D) The same dataset as in (C) fitted with *n*_*c*_ = 2 and *n*_*2*_ = 1.

### Anionic lipids catalyze both primary and secondary nucleation

It is well known that anionic lipids speed up the aggregation of IAPP^20,22–24^. We set out to determine if we could specify which microscopic processes are catalyzed by DOPS. IAPP aggregation assays were performed in the presence of DOPC/DOPS vesicles containing increasing DOPS fractions of 30 %, 50 % and 70 % (**Fig. 5A-C, Fig. S7**). Increasing the DOPS content dramatically increases the speed of aggregation (note the different x-axes in **Fig. 5A-C**), whereas the scaling exponents are similar (**Fig. 5D**). Global fitting of these data (**Fig. 5A-C**) shows that both the combined rate constants for primary and secondary processes, *k*_*+*_*k*_*n*_ and *k*_*+*_*k*_*2*_ (*k*_*+*_ rate constant for elongation, *k*_*n*_ for primary nucleation and *k*_*2*_ for secondary nucleation), scale with increasing DOPS fraction (**Fig. 5E, Table 1**).

**Figure 5.**
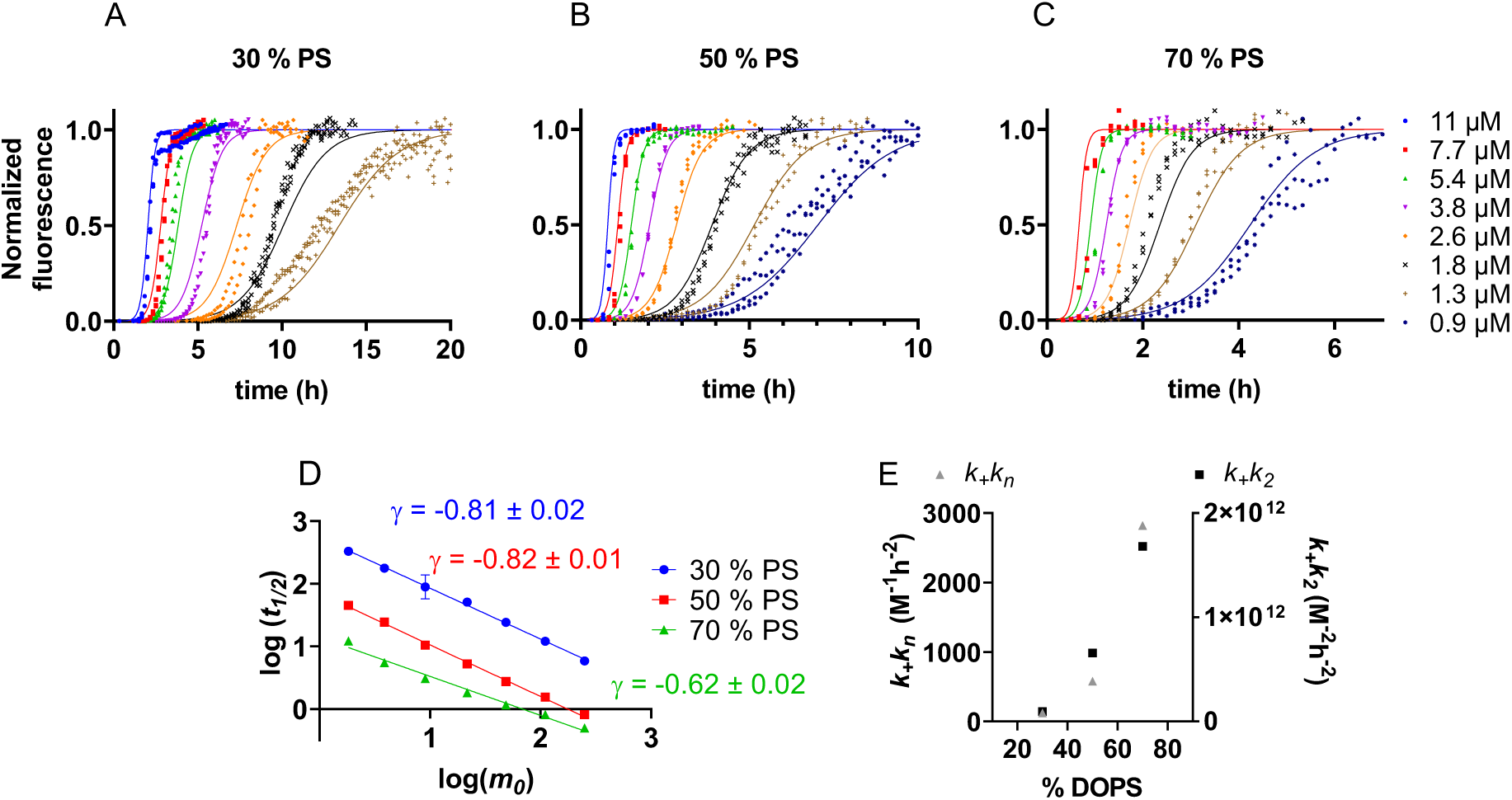
Effect of DOPS on the aggregation kinetics of IAPP. (A-C) Normalized ThT fluorescence data of IAPP aggregation in the presence of DOPC/DOPS vesicles with 30 % (A), 50 % (B) and 70 % (C) DOPS content. Data were fitted to a secondary nucleation dominated model with *n*_*c*_ and *n*_*2*_ of 1. A total IAPP:lipid ratio of 1:50 was used. (D) Scaling exponents of IAPP aggregation in the presence of DOPC/DOPS LUVs with increasing DOPS content. (E) Values of the combined rate constants for primary and secondary processes, *k*_*+*_ *k*_*n*_ and *k*_*+*_ *k*_*2*_, as a function of DOPS content.

The fits to unseeded data do not allow the values for *k*_*+*_*k*_*n*_ and *k*_*+*_*k*_*2*_ to be decomposed into the individual rate constants. Thus, we performed seeded experiments to determine whether DOPS promotes nucleation or elongation. In the presence of fibril seeds, primary nucleation is bypassed, and the increase in fibril mass stems mostly from elongation and secondary processes. At high seed concentrations when many fibril ends are present, elongation dominates at the earliest timepoints, and the initial gradient of the reaction scales with the elongation rate constant *k*_*+*_ (**Materials and Methods, equation 2**).

We confirmed that the addition of fibril seeds strongly reduces the lag phase of IAPP aggregation in the presence of LUVs (**Fig. 6A, B**). Furthermore, the half-times scale linearly with the logarithm of the seed concentration (**Fig. 6C**). These findings are consistent with a mechanism dominated by secondary nucleation^27,29,31^. The initial gradients of highly seeded IAPP aggregation at 30 % and 70 % DOPS content are virtually identical (**Fig. 6D**), revealing that the elongation rate constant is not affected by the DOPS content of the vesicles. Hence, we conclude that the difference in the aggregation kinetics is caused by an increase in the nucleation rates *k*_*n*_ and *k*_*2*_, with increasing DOPS. The data altogether show that the relative abundance of DOPS within the vesicles, but not its total amount increases IAPP nucleation.

**Figure 6.**
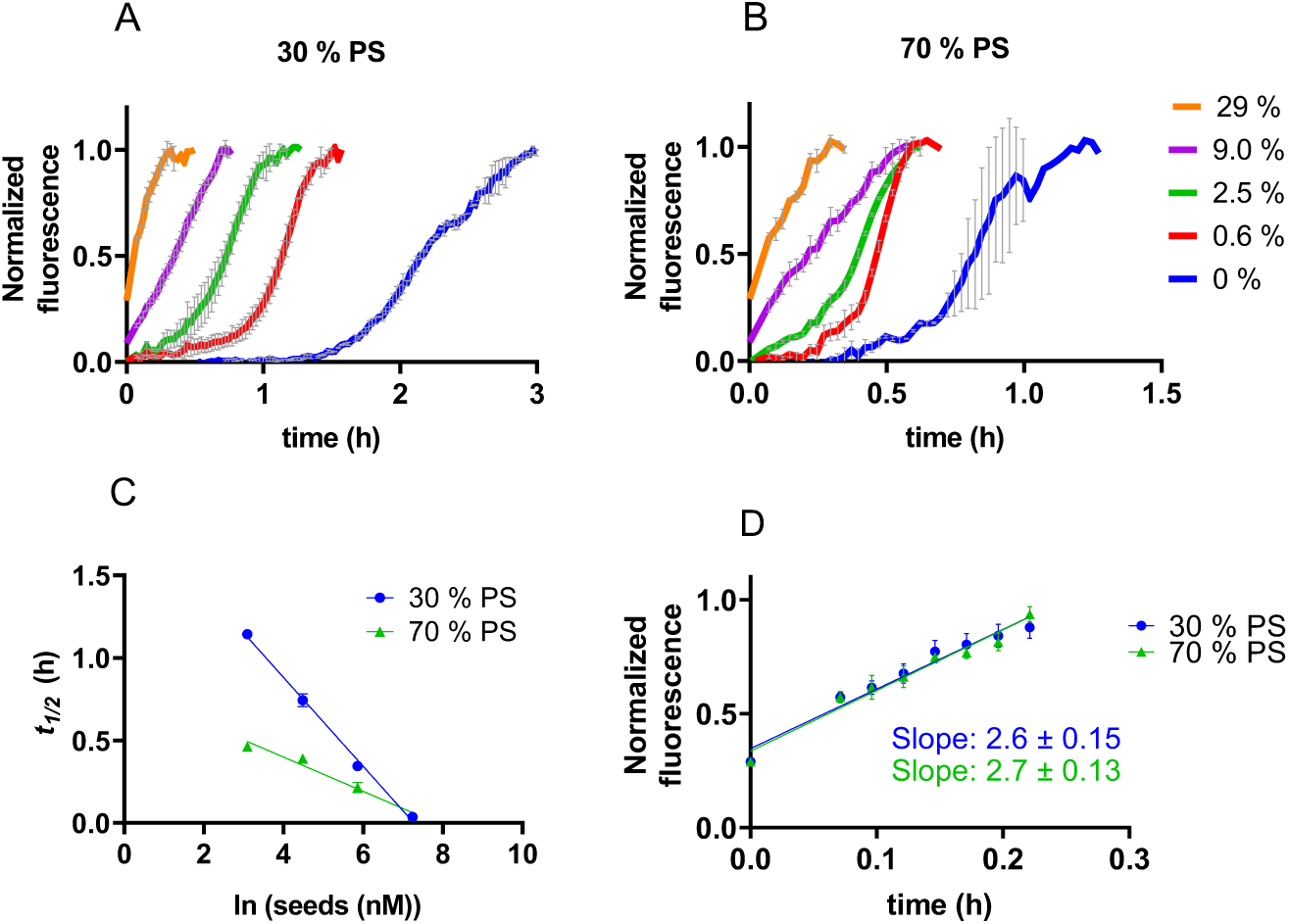
Seeded aggregation of IAPP. (A,B) Normalized ThT-fluorescence of 3.5 µM IAPP with increasing concentrations of seeds with (A) 30 % DOPS or (B) 70 % DOPS LUVs. The seed concentrations are indicated as the percentage of the total IAPP (seeds plus monomers) in monomer units. (C) The logarithm of the seed concentration scales linearly with the half-time of aggregation for 30 % DOPS (blue circles), and 70 % DOPS (green triangles). (D) The initial gradient of a 29% seeded reaction is similar for both DOPS fractions. The data shown in (A), (B), and (C) are the average of triplicates with standard deviation. Seeded experiments were performed at an IAPP monomer:lipid ratio of 1:100.

## Discussion

In conclusion, our data fit to a relatively simple model for IAPP aggregation in the presence of LUVs containing anionic lipids. The key features of the model are: 1) the overall aggregation process is strongly dominated by secondary nucleation, 2) the critical nucleus size comprises a single IAPP monomer, 3) anionic lipids catalyze both primary and secondary nucleation events, but not the elongation process, and 4) these nucleation events thus occur at the membrane surface (**Fig. 7**).

**Figure 7.**
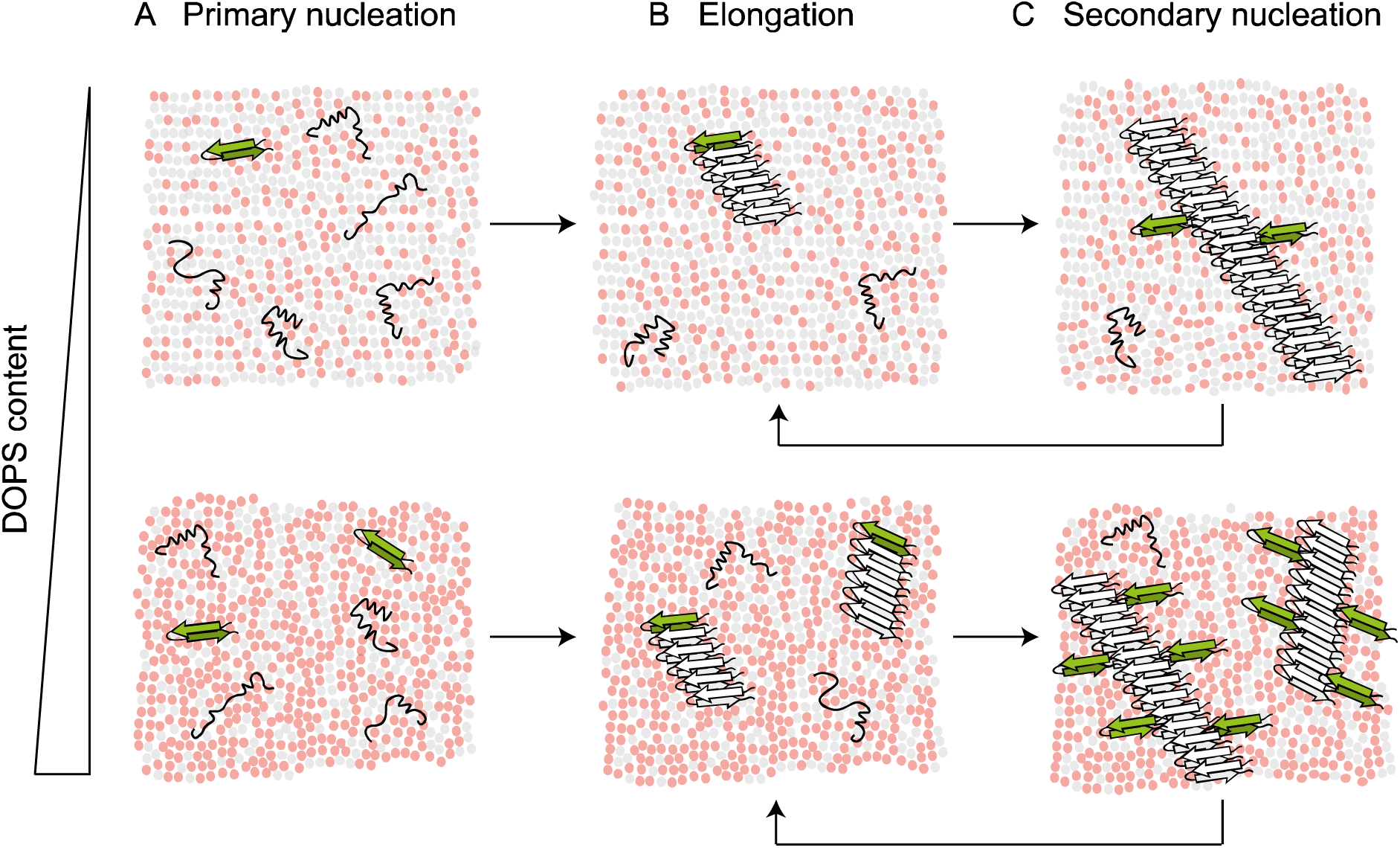
Model for IAPP aggregation catalyzed by membranes containing anionic lipids. (A) IAPP can bind to the membrane in different conformations, some of which may lead to nucleation. Multiple DOPS molecules (red) are involved in the formation of a nucleus (green β-hairpin). (B) Fibril elongation occurs at least to some extent on the membrane surface, but does not depend on the DOPS content. (C) Secondary nucleation occurs at the interface between existing fibrils and DOPS molecules. With increasing DOPS (bottom row), both primary and secondary nucleation are more strongly catalyzed.

Despite the fact that secondary nucleation has also been reported to dominate IAPP aggregation in solution^14–16^, the aggregation mechanism on lipid membranes is fundamentally different given that it is restricted to a two-dimensional surface. The reaction order for nucleation of IAPP in solution was found to be closer to 2^16^. Although the IAPP used in this study was prepared in a different manner, the comparison suggests that the monomeric IAPP nucleus we observe here may be the result of a specific conformation coordinated by DOPS molecules. The conversion of pre-existing oligomers into nuclei would also result in a reaction order of 1, and we cannot formally exclude this possibility. However, IAPP has previously been reported to insert as a monomer into membranes with the same composition as used here^18^, and our data argue against a subsequent assembly into an oligomeric nucleus.

The possibility of a monomeric nucleus is supported by the observation that the aggregation mechanisms and rates do not depend on the local density of IAPP on the membrane. The aggregation kinetics remain approximately the same over a range of peptide:lipid ratios of 1:10 to 1:100 (**Fig. 2, Fig. 3, Table 1**), which effectively corresponds to a ten-fold dilution of IAPP on the membrane. When fitting datasets obtained at a fixed peptide:lipid ratio (**Fig. 2, Fig. 3**), the peptide density on the vesicles is expected to be identical for all IAPP monomer concentrations, and processes depending on the local crowding on the membrane surface are not taken into account. However, these processes do not appear to play a role. Indeed, aggregation data obtained at a constant concentration of lipids, leading to increased crowding of IAPP on the membrane with increasing monomer concentration, fit to the same mechanism with a reaction order for primary and secondary nucleation of 1 (**Fig. 4**).

Increasing the fraction of DOPS in the vesicles strongly accelerates the aggregation kinetics of IAPP (**Fig. 5**), in agreement with stronger binding of IAPP at higher negative charge of the membrane. It is conceivable that multiple DOPS molecules are involved in the nucleation of one IAPP monomer, which would lead to a more than linear increase in the nucleation rates with increasing DOPS fraction. Interestingly, DOPS speeds up both primary and secondary nucleation of IAPP, which is a novel mechanism that has so far not been reported for other lipid and protein systems. Only primary nucleation was accelerated for amyloid-β in the presence of LUVs containing cholesterol, whereas the rate constant of secondary nucleation was unaffected^32^. Also for α-synuclein, only primary nucleation was promoted by purely anionic DMPS vesicles, in the absence of secondary processes^33^.

The finding that DOPS promotes the secondary nucleation of IAPP implies that fibril elongation occurs at least in part on the membrane surface (**Fig. 7**). This conclusion is in agreement with previous findings for IAPP aggregation, where fibril elongation was implied as a cause of vesicle leakage^12^. Our data raise the possibility that secondary nucleation of IAPP monomers may also contribute to membrane damage, as these events take place over the same time course. Whereas we cannot exclude the formation of oligomeric species, they do not appear essential to explain membrane-catalyzed IAPP aggregation according to the model that we here establish.

Altogether, our results reveal a previously undescribed molecular mechanism for IAPP aggregation catalyzed by lipid membranes, and explain the role of anionic lipids in this process. This framework will be important for future studies on the mechanisms by which IAPP damages cellular membranes and causes β-cell death.

## Associated content

The following files are available free of charge.

Supporting information containing raw ThT data and additional fits (PDF).

## Author information

### Author Contributions

The manuscript was written through contributions of all authors.

