## Supplementary Information for "Membrane-catalyzed aggregation of islet amyloid polypeptide is dominated by secondary nucleation"

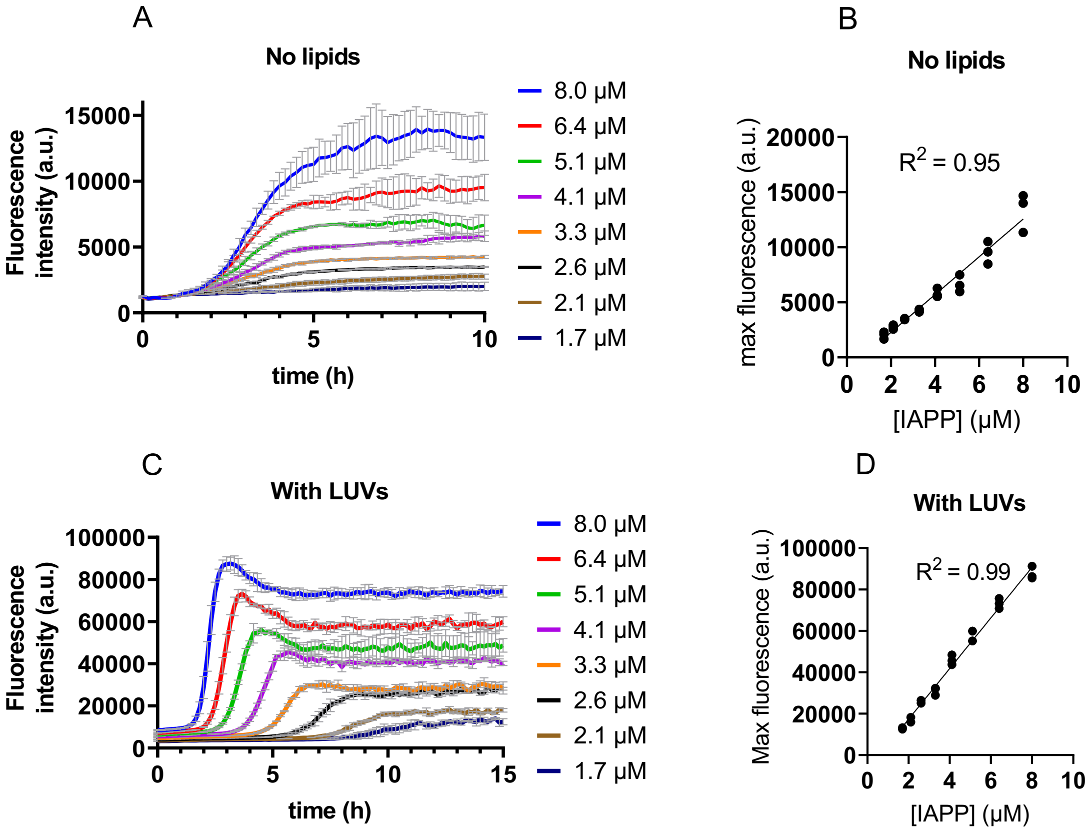


**Figure S1**. Raw fluorescence data accompanying Figure 1. (A, C) Raw ThT fluorescence data of IAPP aggregation in the absence of lipids (A), or in the presence of LUVs at 1:100 IAPP:lipid molar ratio (C). The curves show the average of triplicates with standard deviation. (B, D) Correlation between the fluorescence at the plateau and the concentration of IAPP for the respective datasets on the left. 10 μM of ThT was found to be optimal to monitor the aggregation of IAPP in these experiments.

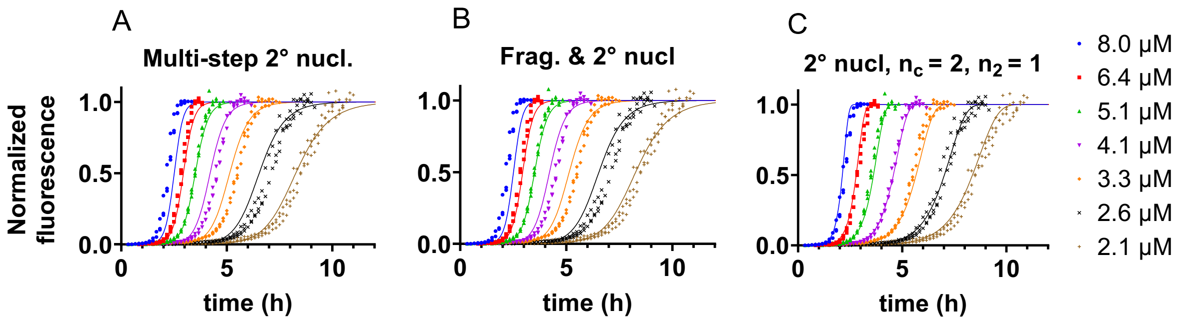


**Figure S2.** Additional fits for IAPP aggregation kinetics at 1:100 peptide:lipid ratio. (A) Fits to a multi-step secondary nucleation mechanism ($n_c$ and $n_2$ = 2). (B) Fits to a mechanism that includes both fragmentation and secondary nucleation ($n_c$ and $n_2$ = 2). (C) Fits to a model dominated by secondary nucleation with a reaction order for primary nucleation ($n_c$) of 2, and a reaction order for secondary nucleation ($n_2$) of 1.

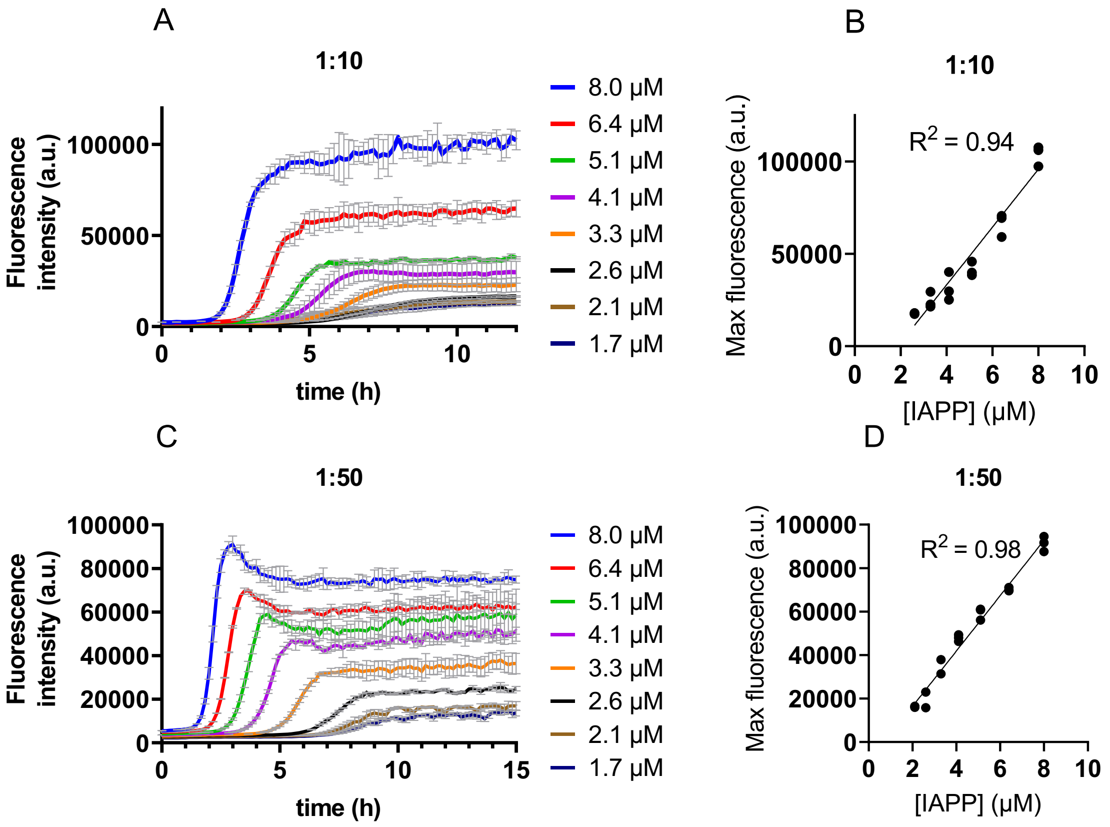


**Figure S3.** Raw ThT data and scaling exponents of IAPP at 1:10 and 1:50 peptide:lipid ratios. (A, C) Raw ThT fluorescence data of IAPP aggregation in the presence of LUVs at 1:10 (A) or 1:50 (C) IAPP:lipid molar ratio. The curves show the average of triplicates with standard deviation. (B, D) Correlation between the fluorescence at the plateau and the concentration of IAPP for the respective datasets on the left.

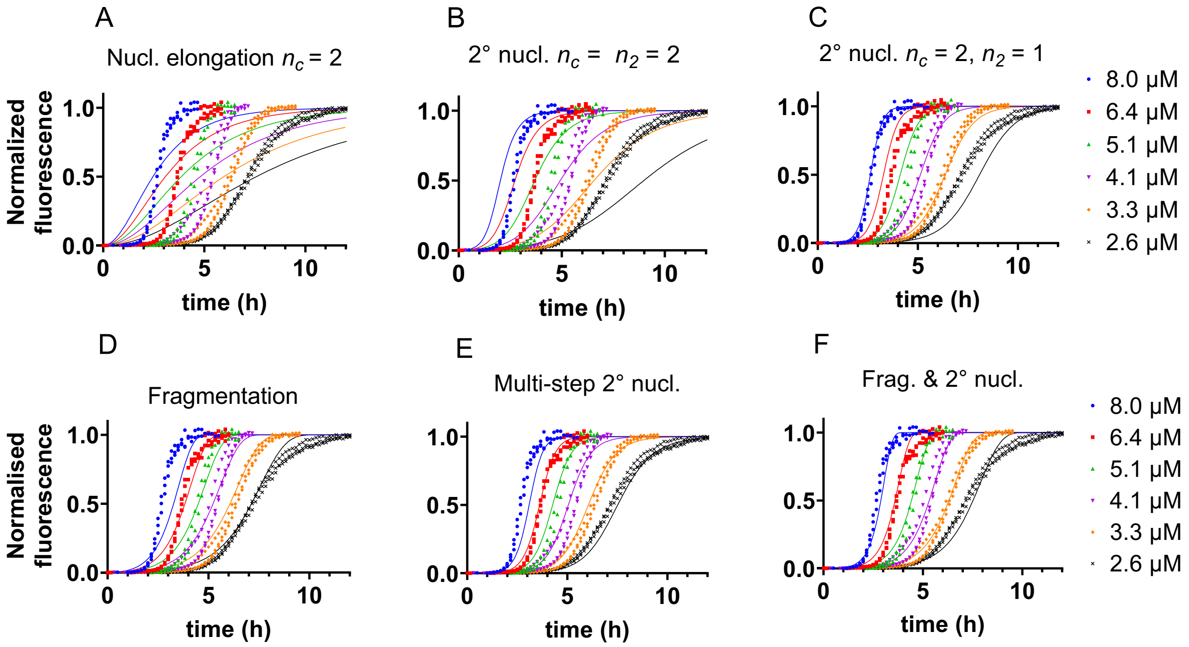


**Figure S4.** Additional global fitting of IAPP aggregation kinetics at 1:10 peptide:lipid ratio. (A) Fits to a model of nucleation and elongation with a reaction order for nucleation ($n_c$) of 2. (B) Fits to a model dominated by secondary nucleation with reaction orders for primary nucleation ($n_c$) and secondary nucleation ($n_2$) of 2. (C) Fits to a model dominated by secondary nucleation with a reaction order for primary nucleation ($n_c$) of 2, and a reaction order for secondary nucleation ($n_2$) of 1. (D) Fits to a model dominated by fragmentation. (E) Fits to a multistep secondary nucleation mechanism ($n_c$ and $n_2 = 2$). (F) Fits to a mechanism that includes both fragmentation and secondary nucleation ($n_c$ and $n_2 = 2$).

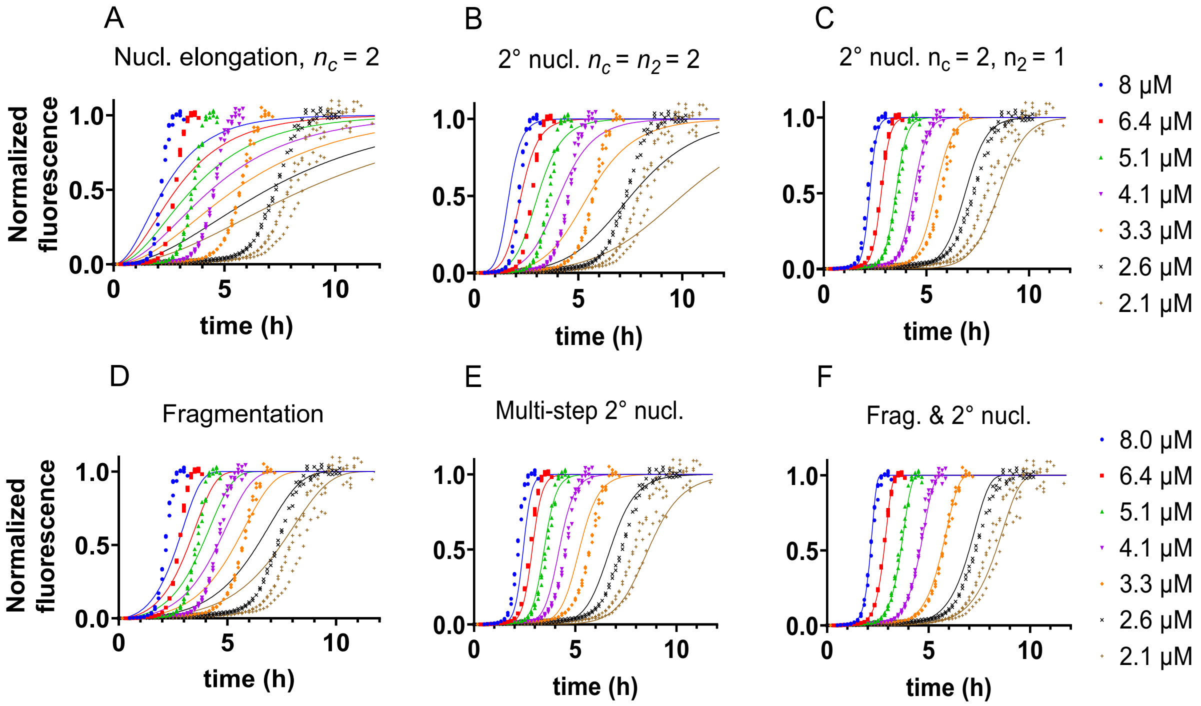


**Figure S5.** Additional global fitting of IAPP aggregation kinetics at 1:50 peptide:lipid ratio. (A) Fits to a model of nucleation and elongation with a reaction order for nucleation ($n_c$) of 2. (B) Fits to a model dominated by secondary nucleation with reaction orders for primary nucleation ($n_c$) and secondary nucleation ($n_2$) of 2. (C) Fits to a model dominated by secondary nucleation with a reaction order for primary nucleation ($n_c$) of 2, and a reaction order for secondary nucleation ($n_2$) of 1. (D) Fits to a model dominated by fragmentation. (E) Fits to a multistep secondary nucleation mechanism ($n_c$ and $n_2 = 2$). (F) Fits to a mechanism that includes both fragmentation and secondary nucleation ($n_c$ and $n_2 = 2$).

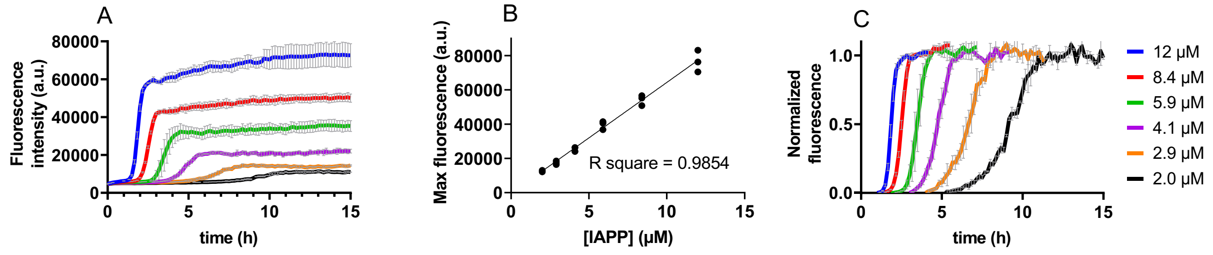


**Figure S6.** Raw and normalized data for IAPP aggregation with a constant lipid concentration of 900 µM. (A) Raw ThT fluorescence data for IAPP aggregation in the presence of 900 µM LUVs. (B) Correlation between the fluorescence at the plateau and the respective IAPP concentration. (C) Normalized aggregation curves of the data shown in (A). The curves show the average of triplicates with standard deviation.

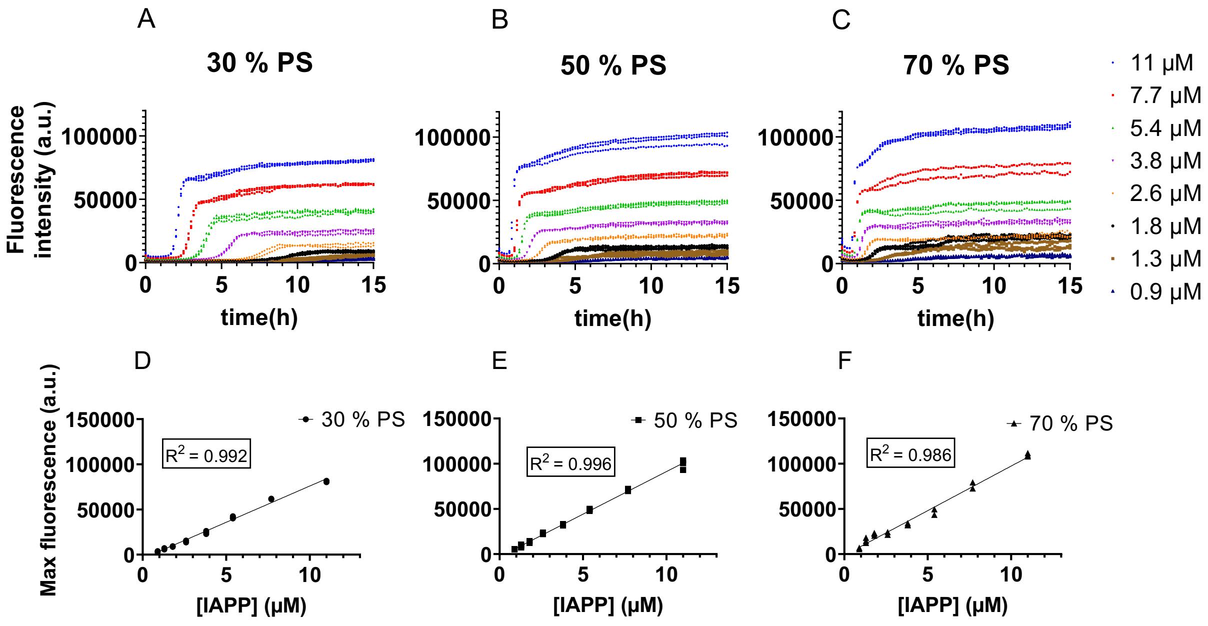


**Figure S7.** Raw and normalized data for IAPP aggregation at different DOPS fractions. (A, B, C) Raw ThT fluorescence data for IAPP aggregation in the presence of 1:50 LUVs containing 30 % (A), 50 % (B), or 70 % (C) DOPS. (D, E, F) Correlation between the fluorescence at the plateau and the respective IAPP concentration.
